# Riemannian Geometry Boosts Representational Similarity Analyses of Dense Neural Time Series

**DOI:** 10.1101/232710

**Authors:** Alexandre Barachant, Jean-Rémi King

**Affiliations:** New York University (USA); Frankfurt Institute for Advanced Studies (Germany)

**Keywords:** MEG, Decoding, Representational Similarity Analyses, Riemannian Geometry, multivariate pattern analysis

## Abstract

Representational similarity analysis (RSA) is a popular technique to estimate the structure of mental representations from neuroimaging data. However, RSA can be difficult to estimate for neural time series, where mental representations may be distributed in a highly dimensional space. Here, we show that RSA can be efficiently estimated from dense neural time series using Riemannian geometry. Using a public magneto-encephalography dataset, we decoded 24 classes from the brain evoked responses to 720 visual stimuli. RSA estimated from the confusion matrices of a standard regularized logistic regression achieved an average decoding accuracy of 23% (chance=4%). Our approach based on spatial filtering and Riemannian geometry nearly doubled this score with an average 42% decoding accuracy. Finally, our results revealed how RSA becomes ill-conceived when it derives from confusion matrices of highly accurate multivariate pattern classifications. Instead, we propose to directly estimate RSA from Riemannian metrics without fitting a multivariate pattern classifier. Overall, our approach, based on Riemannian geometry provides a principled and efficient basis to study the structure of mental representations from highly dimensional neural time series.

## Introduction

Representational similarity analyses (RSA) is an increasingly popular technique to estimate the structure of mental representations from brain activity (Kriegeskorte 2008). It consists in 1) cross-correlating the multidimensional brain responses to many stimuli recorded in a given area (e.g. Kriegeskorte 2008) and/or at a given time sample (e.g. Cichy et al. 2014) and 2) assessing the clustering from the resulting cross-similarities of neural patterns. In practice, this ‘dissimilarity matrix’ is estimated either from the correlation of neural patterns across conditions across (a form of Euclidian distance across neural patterns) or from the confusion matrices of one-versus-all linear classifiers trained to predict the stimulus class from the neuroimaging data (e.g. Cichy et al. 2014).

This approach to RSA poses two challenges. First, it does not trivially extend to temporally-resolved neuroimaging data, where neural contents may be represented over multiple time samples: time can rapidly increase dimensionality which leads Euclidian metrics to poor estimates of pattern similarities. Second, confusion matrices are only informative in medium signal-to-noise ratio (SNR) conditions (Fig. 1). Indeed, low SNR prevents decoding performance, and thus lead to random confusion matrices, whereas perfect SNR ratio can lead to strictly diagonal matrices, where no confusion is made across stimuli independently of their true clustering.

**Figure 1.**
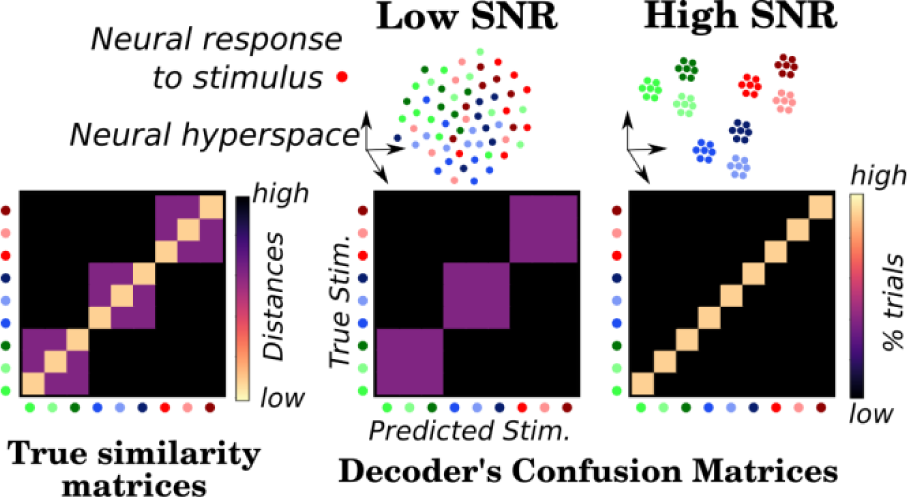
The confusion matrices of classifiers trained to decode hyperdimensional neural responses can confound signal-to-noise ratio (SNR) and the geometry of neural representations.

Here, we propose a framework to perform RSA that addresses these two issues. It capitalizes on the notion that dense neural time series can be analyzed in terms of covariance matrices, which, under normality assumption, correspond to unique locations on a Riemannian manifold (Barachant et al 2012-2013). The Riemannian distances between each set of temporally-resolved brain responses can be directly estimated, and provide an efficient proxy to estimate the clustering of neural representations.

## Methods

We used the public dataset from Cichy et al (2014) as provided in *MNE*. It is composed of four runs of a single subject recorded with a 306-channel Neuromag Elekta magneto-encephalography (MEG) sampled at 1 kHz, while being presented with static images. Here we focus on 720 trials from the first 24 classes (12 faces and 12 body parts). Raw MEG data were epoched from 0 to 600 ms after stimulus onset, baseline-corrected between −100 and 0 ms and decimated by four. Two multivariate approaches were implemented, each ending with an l2-regularized (C=1) logistic regression (LR) classifier. The first ‘Euclidian’ approach consisted in vectorizing the MEG data (each trial being a 306 MEG channels * 150 time samples vector). The second, ‘Riemannian’ approach consisted in applying 2 Xdawn spatial filters (Rivet et al, 2009), and projecting the resulting event related field meta-covariance data (Congedo 2017) into the tangent space with *pyRiemann*. For both approaches, we estimated the distances between neural patterns as well as the confusion matrices of the logistic regression (the % of trials predictions for each class). All analyses were performed at the sensor level using evoked responses with an 8-split stratified shuffle-split cross-validation procedure with *Scikit-Learn*.

## Results

The LR fit with Euclidian data led an average accuracy of 23.3% (±2% standard error of the mean [SEM] across categories, chance=4%); Computation time took 23 minutes on an 8-core laptop. The Riemannian classifier led to an average 41.3 % (±4%) computed in 16 minutes (Fig 2. Top). Our results demonstrate an important SNR gain in using a spatial filtering with Riemannian geometry. However, decoding performance is so high that the corresponding confusion matrix may be inappropriate for estimating the clustering of neural responses (Fig. 1.). This limitation is largely overcome when using distance metrics (Fig 2. Bottom). Specifically, the Riemannian distances both maximize SNR and reveal the clustering of neural responses to face stimuli.

**Figure 2:**
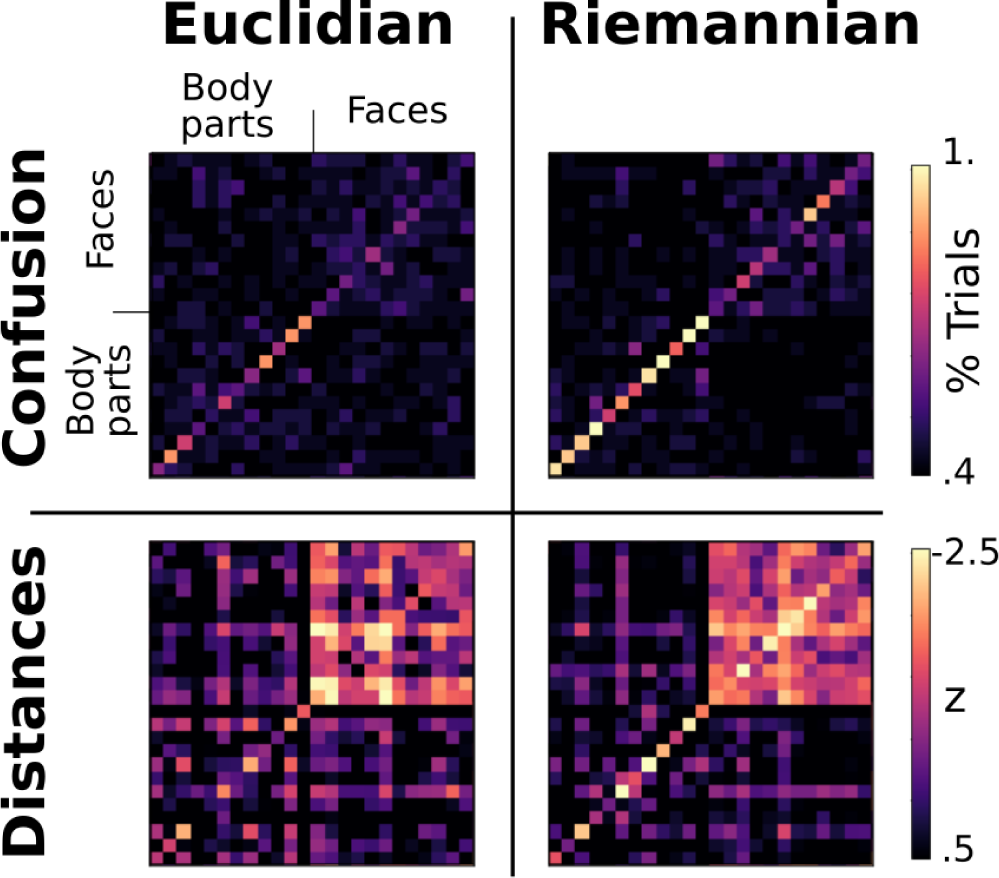
Top. The confusion matrices for the two approaches. Bottom. the corresponding distances, z-scored for comparison purposes.

## Conclusions

Our approach shows how RSA can be applied to high dimensional neural time series by using Riemannian geometry. Our results both maximize SNR and allow a direct assessment of the similarities of neural responses. The present approach thus holds great promises to investigate the structure of mental representations from dynamic neural activity.

## Acknowledgments

European Union’s Horizon 2020 research & innovation program under the Marie Sklodowska-Curie Grant Agreement No. 660086, Bettencourt-Schueller and the Philippe Foundations (J-R.K).

